# An 8-cage imaging system for automated analyses of mouse behavior

**DOI:** 10.1101/2023.02.04.527129

**Authors:** Thaís Del Rosario Hernández, Narendra R. Joshi, Sayali V. Gore, Jill A. Kreiling, Robbert Creton

## Abstract

The analysis of mouse behavior is used in biomedical research to study brain function in health and disease. Well-established rapid assays allow for high-throughput analyses of behavior but have several drawbacks, including measurements of daytime behaviors in a nocturnal animal, effects of animal handling, and the lack of an acclimation period in the testing apparatus. We developed a novel 8-cage imaging system, with animated visual stimuli, for automated analyses of mouse behavior in 22-hour overnight recordings. Software for image analysis was developed in two open-source programs, ImageJ and DeepLabCut. The imaging system was used to measure multiple behaviors, including acclimation to the novel cage environment, day and nighttime activity, stretch-attend postures, location in various cage areas, and habituation to animated visual stimuli. These behaviors were summarized in behavioral profiles, which may be used in further studies to examine treatments for neural disorders.

## Introduction

Rapid tests are available for mouse behavioral phenotyping (*1*). Behavioral tests such as the open-field test and elevated plus maze have been used for decades and are still widely used to examine mutant lines, transgenic lines, and therapeutic treatments (*2–4*). Many rapid tests can be carried out within 15 minutes, which is critical when examining a large number of mice in a high-throughput setting. However, these short tests have several drawbacks. 1) The nocturnal mice are typically imaged in a well-lit environment at daytime when mice normally hide and sleep. 2) Pre-test handling is a major confounding factor, as behaviors are evaluated immediately after handling. 3) Behaviors are measured without acclimation to the testing apparatus, and 4) mice need to be moved between different testing devices for the analysis of multiple behaviors. While some of these drawbacks can be mitigated by standardized procedures and attention to experimental detail, questions have been raised about the robustness of behavioral assays and the repeatability of the obtained results (*5–7*).

Home-cage imaging may be used as an alternative approach to measure behavior over several days (*7*). The extended imaging period contributes to the acquisition of robust results and can reveal additional behavioral phenotypes, such as changes in circadian rhythms. At the forefront of this rapidly expanding field are novel image-analysis techniques to recognize animal postures in a complex background (*8–11*). However, it can be challenging to image large numbers of mice for days on end.

The current study describes the development of an 8-cage imaging system with a projector to examine responses to visual stimuli during 22-hour overnight recordings. This imaging system fills the gap between rapid tests and home-cage analyses. A single imaging system can examine multiple behaviors in 32 mice per week. In addition, the overnight recordings allow for analyses of behavior during daytime and nighttime, for an extended period after animal handling, both during and after acclimation to the novel cage environment.

## Results

### The 8-cage imaging system

We built an 8-cage imaging system for automated analysis of mouse behavior (Fig 1). The imaging system was designed to be inexpensive, easy to use, and suitable for the analysis of multiple behaviors in an overnight recording. The system was built in a transmitted light configuration, with two cameras above the cages and a projector below the cages. The projector provides background illumination during daytime (white light) and nighttime (red light), using the white opaque bottom of the cages as a back-illuminated screen. PowerPoint presentations with automated transitions were used to switch from daytime to nighttime and to display visual stimuli at nighttime when the nocturnal mice are most active (Supplementary information, S1). We found that this imaging system is capable of acquiring high-contrast images (Fig 2). All mice are visible throughout the imaging experiment, even when mice are hiding in their transparent red houses and when using red background illumination. The mouse cages have different environments in each quadrant of the cage, providing enrichment and extending the range of behaviors that can be measured. The ‘Home’ area has a red mouse house for shelter. The ‘Wall’ area provides an open field surrounded by two plain cage walls. The ‘Food’ area has a food and water tray, and the ‘Window’ area has a screened in window. This window is used for ventilation and to give a view of another mouse outside the window. The two camera feeds were combined in a single 22-hour movie, stored at 1 frame per second in a compressed AVI format. These 9 gigabyte (GB) movies can be played back at 30x speed (under an hour) to provide an overview of the imaging experiment. The 22-hour movies of 8 mice contain a wealth of information, suitable for automated image analysis.

**Fig 1.**
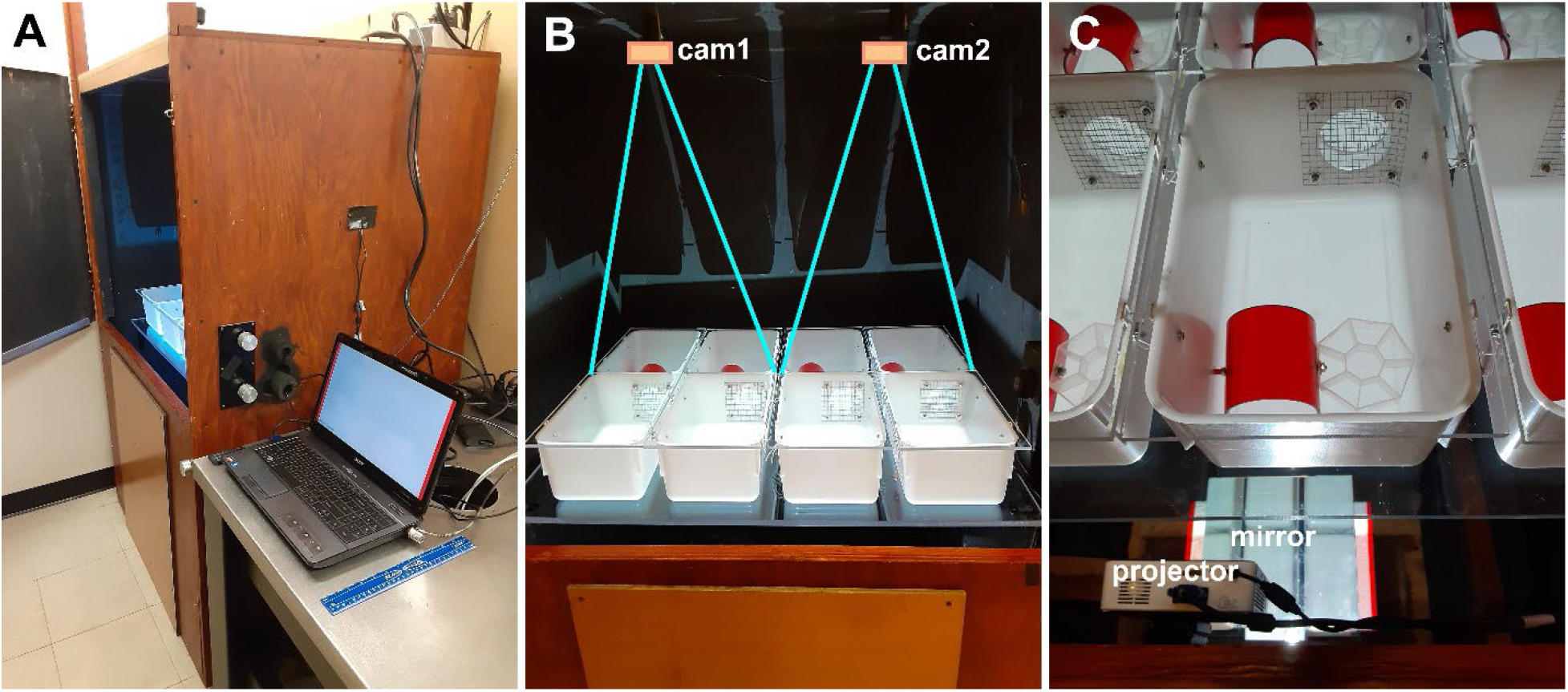
The 8-cage imaging system. A) Dark box and computer. B) Eight cages on a transparent stage, imaged with two cameras, cam1 and cam2. C) A projector illuminates the bottom of the cages via a mirror that lays flat on the floor. The cages contain a red transparent home, an 8-compartment food/water tray, and a screened window. Acrylic lids cover the cages. The top of the cage is 18.5 cm wide and 29.4 cm long. The red homes are 7.3 cm wide and 7.6 cm long.

**Fig 2.**
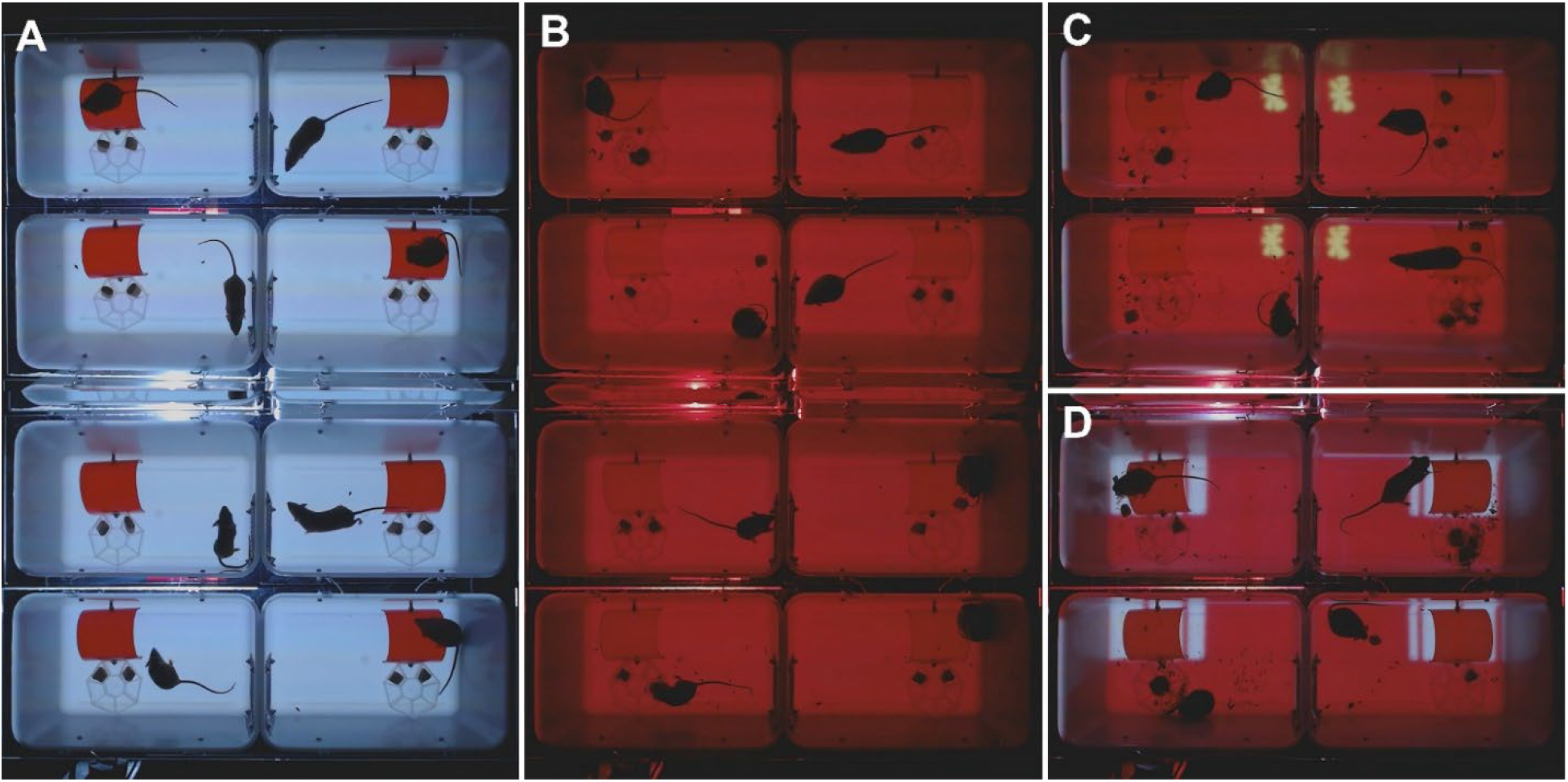
Acquired images on the 8-mouse imaging system. A) Daytime with white lighting. B) Nighttime with red lighting. C) Yellow moth in wall quadrant. D) Moving lines in home quadrant. The red homes are 7.6 cm long.

### Image analysis in ImageJ and Fiji

ImageJ and Fiji are two freely available image analysis programs, which allow for complex analyses with a point-and-click interface. We developed an ImageJ macro (Supplementary information, S2), which can be loaded as a plugin in either ImageJ or Fiji (Fig 3). The macro has a dialog boxes to guide the user in opening a virtual stack, defining regions of interest, and calibrating the camera timer. It then takes the red channel of an RGB image, selects a cage, applies a threshold, and carries out a particle analysis. The particle analysis filters out smaller particles (food) and measures the centroid, bounding box, and best-fitting ellipse of the mouse. This process is automatically repeated for all cages in an image and for all images in the virtual stack. The measurements are saved in a ‘Results’ file with a long list of X and Y coordinates. The macro then calculates additional parameters, such as position of the head, speed, movement, location in the cage, and the stretch-attend posture (SAP). For quality control, the macro shows the measurements on the first image (green dot for centroid, red dot for calculated head region). This quality control measure revealed that the centroid of the mouse can be reliably identified. However, the head region can be miscalculated when a mouse curls up. Based on this initial finding, we adjusted the macro to only label the head region when a mouse is stretched out. In addition, we did not use the head position for the subsequent analyses of behavior.

**Fig 3.**
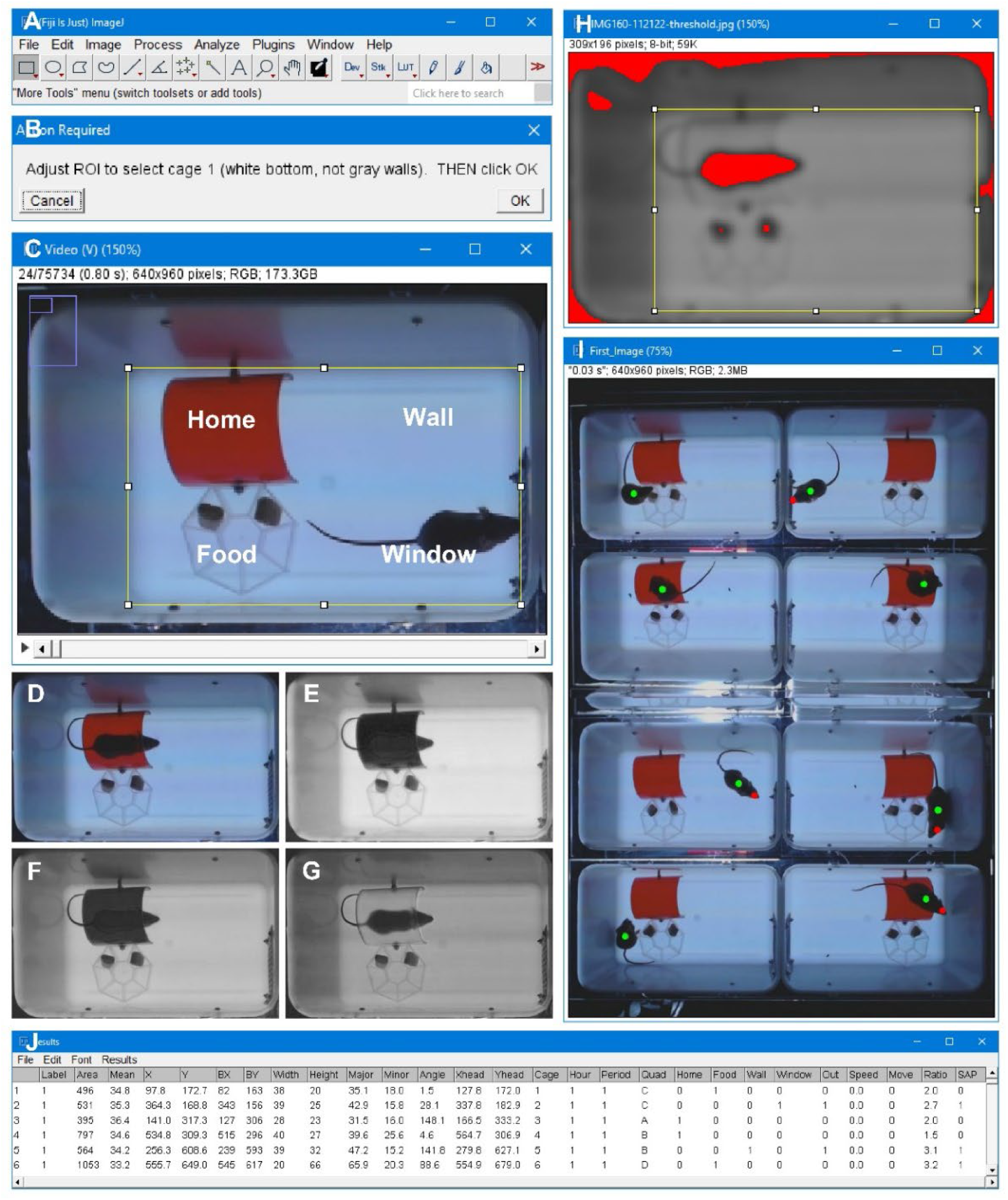
ImageJ analysis. A) The analysis can be carried out in ImageJ or Fiji. B) A custom-developed macro guides the user in setting up eight regions of interest (ROIs). C) Region of interest, which sets four equal-sized quadrants in the cage; home, wall, food, and window. D) Full-color image with a mouse in a red house. The macro splits this color image into three gray-scale channels: E) a blue channel, F) a green channel and G) a red channel, which is used for further analysis (the mouse is visible in the red channel). H) The image is blurred and a threshold is applied to separate dark objects from the background. Objects are identified by particle analysis, which excludes objects smaller than 300 pixels. This process is automatically repeated for all images in a video. I) The macro displays the measurements on the first image for quality control. J) Data is logged in a ‘Results’ file, which shows both measured values (e.g. X, Y of centroid) and calculated values (e.g. location of a mouse in a quadrant).

### Image analysis in DeepLabCut

DeepLabCut (DLC) is an open-source program for automated analyses of behavior. This python-based program uses training sessions and artificial intelligence to recognize specific points of interest in a complex background. We examined if this program can be used as an alternative to ImageJ and Fiji for the automated analysis of behavior in our 8-cage imaging system (Fig 4). DLC was used to train a network using various videos of mice in the 8-cage imaging system. In total 430 frames were extracted and labeled with 48 points of interest: (1) nose, (2) right ear, (3) left ear, (4) head, (5) neck, (6) center, (7) hip, (8) tail base, (9) right front paw, (10) left front paw, (11) right rear paw, (12) left rear paw, (13) food plate, (14) food pellet 1, (15) food pellet 2, (16) front-top house, (17) front-bottom house, (18) back-top house, (19) back-bottom house, (20) center house, (21) center-back house, (22) up-moth top, (23) up-moth bottom, (24) up-moth right wing, (25) up-moth left wing, (26) down-moth top, (27) down-moth bottom, (28) down-moth right wing, (29) down-moth left wing, (30) up-worm head, (31) up-worm center, (32) up-worm tail, (33) down-worm head, (34) down-worm center, (35) down-worm tail, (36) black cricket, (37) black swoop, (38) black loom, (39) yellow cricket, (40) yellow swoop, (41) grey loom, (42) flash, (43) cage top left, (44) cage top right, (45) cage bottom left, (46) cage bottom right, (47) window top, (48) window bottom. These points were chosen to obtain a wide range of mouse behaviors, including interactions with objects in the cage (e.g. hut, food pellets) as well as distance from specific stimuli regions, even when those stimuli move around the cage. Multiple stimuli are included in the model even when the video to be analyzed may not have an instance of all stimuli. This allows for flexibility of stimuli presentation without having to train a new model. We recorded 8 mice in the 8-cage set-up for a total of 22 hours and, to allow the DLC model to perform inference for each individual mouse, the video was cropped around the edges of the home cage. We used a minimum and maximum area parameters to accurately identify the cropping coordinates for each cage. The code implementation used for cage identification and cropping can be found in our source code on GitHub (https://github.com/Creton-Lab). Our trained DLC model recognizes the points of interest in different lighting scenarios, including both “day” (white light) and “night” (red light), and with a paper towel, which some mice use to build a nest.

**Fig 4.**
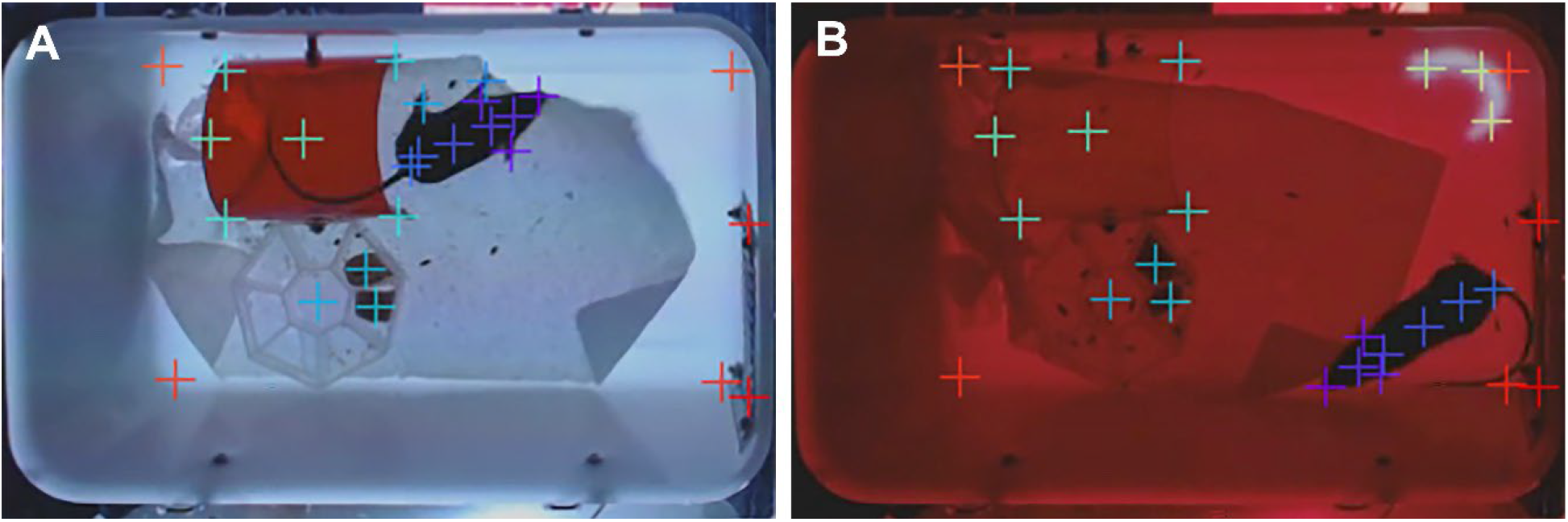
Analysis in DeepLabCut. A) Specific structures are identified after training a model in DeepLabCut. The current model various parts of the mouse, cage, and visual stimuli. B) These structures can also be identified at nighttime, using red light for background illumination, with or without nesting materials and visual stimuli.

### Optimization of the imaging system with two mice

The imaging system, cages, and methods for image analysis were optimized using two mice (Fig 5). Both were female B6129SF2/J mice with a mixed genetic background. In contrast to inbred lines, each B6129SF2/J mouse is genetically unique. The two mice were easily recognized; one mouse had black fur and the other mouse had agouti fur. The two mice were imaged repeatedly (once or twice a week), using various cage configurations and visual stimuli. The mice were imaged with a single mouse house (red) or two mouse houses (red and amber) and were imaged in different cages, with and without nesting material (a paper towel). All experiments started with 6 hours of daytime and 6 hours of nighttime, followed by various visual stimuli, including moving red lines on a white background in the home area, a circling gray worm, a rotating yellow moth, multiple yellow moths, and a moving cricket. When watching the acquired movies at 30x speed, the mice seemed most active during the first hour of imaging (acclimation period), least active in the following 5 hours (daytime), moderately active at nighttime, and only transiently active in response to novel visual stimuli, such as the rotating yellow moth and moving lines in the home area. In all imaging experiments, the black mouse was more active and exploratory than the agouti mouse. The black mouse frequently moved between the home and window area. It would build a nest in its home when nesting materials were available, and in some experiments the nest was so dense that it would interfere with the automated image analysis. In contrast, the agouti mouse would stay in its home for extended periods, would not build a nest, and would only briefly explore other areas in the cage. While these 2-mice experiments were not suitable for statistical analyses, we did use the results to fine-tune the imaging configuration, visual stimuli, and analysis. We decided to move forward without nesting materials, using the yellow moth and moving lines as visual stimuli. Acclimation to the imaging system was measured by comparing the 1^st^ vs. 2^nd^ hour of imaging. Habituation to visual stimuli was measured by comparing activity during the first 10 minutes vs. the next 50 minutes. The consistent differences between the black and agouti mouse in multiple experiments suggest that the 8-cage imaging system can be used to distinguish genetically unique mice, irrespective of external factors such as handling, availability of nesting materials, cage position, and repeated measurements over time.

**Fig 5.**
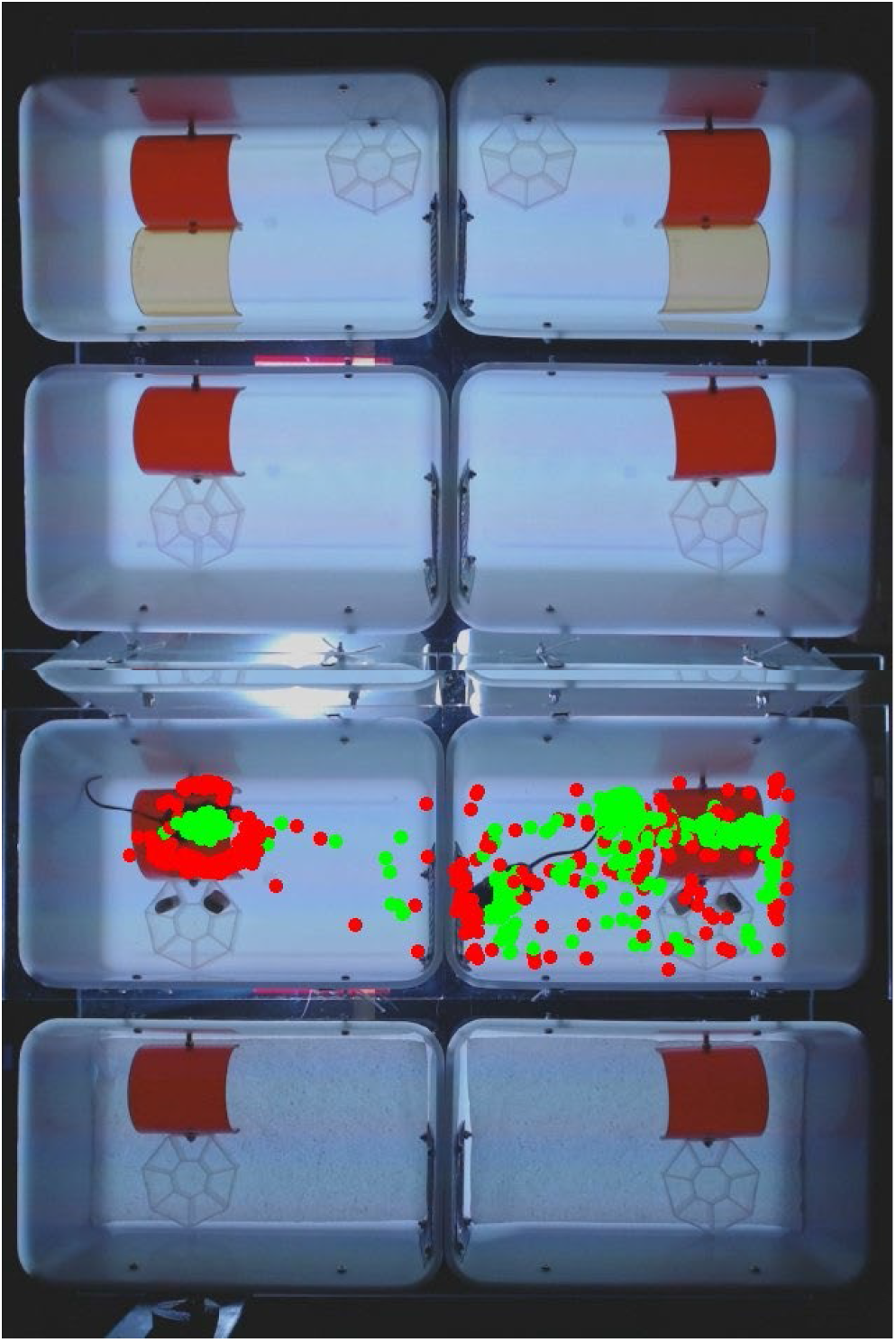
Optimization of imaging parameters. The imaging parameters were optimized using two mice, an agouti mouse (left) and black mouse (right), which were imaged in five separate experiments. In each experiment, the agouti mouse displayed low activity and preferred to stay in its house (cage 5), while the black mouse was more active and exploratory (cage 6). Top two cages (cage 1 and 2): red and amber mouse house. Center four cages: red mouse house. Bottom two cages (cage 7 and 8): red mouse house with paper towel. Green dots = centroid of mouse in the first 600 images. Red dots = head of mouse in first 600 images (10 min). Mouse house = 7.6 cm long.

### Analysis of behavior in 8 mice

The developed imaging system was used to image 8 female B6129SF2/J mice that had not been imaged before. We imaged these mice for 22 hours and used the ImageJ macro for the analysis of behaviors. The ImageJ ‘Results’ file was copied into a MS Excel template (Supplementary information, S3), which then summarizes the obtained data in tables and graphs (Supplementary information, S4). The datasets in S4 are provided in a user-friendly format that is well suited for further data mining, i.e. researchers can examine various behaviors in MS Excel. We focused our studies on activity, the stretch-attend posture, and time spent in each quadrant of the cage. These behaviors are described in more detail below.

### Analyses of activity

Changes in activity over time were examined in 22-hour graphs (Fig 6). Activity was measured as speed (in cm / sec) and the percentage of time that the mice move more than 1 cm / sec (% move). While the individual mice displayed considerable variability, clear patterns emerged when averaging measurements of 8 mice (Fig 6A-D). Specifically, the mice displayed a higher average speed during the first hour of imaging (0.97 cm/sec) than the second hour of imaging (0.30 cm/sec), and displayed a higher average speed at nighttime (0.70 cm/sec) than at daytime (0.28 cm/sec). These difference were significant (Welch test, p=2×10^−4^ and p=3×10^−3^, respectively). Similar results were obtained when measuring the percentage of time that the mice move, i.e. the mice move more in the first hour (19.7%) than the second hour (6.3%) and move more at nighttime (12.7%) than at daytime (5.1%). These differences were again significant (Welch test, p=3×10^−4^ and p=9×10^−4^ respectively). The difference in movement between these the first and second hour (13%) may be used as a measure of acclimation. To analyze responses to visual stimuli, the percentage of time that the mice move was analyzed in 10 minute periods (Fig 6E). The rotating yellow moth induced a 10% increase in activity during the first 10 minutes and 10% decrease in activity afterwards, but neither effect was statistically significant. The first set of moving lines (15^th^ hour, period 85-90) induced a 10% increase in activity during the first 10 minutes (not significant), and a 16% reduction in activity afterwards (Welch test, p=4×10^−3^), suggesting that the mice habituate to the moving lines after 10 minutes. The second set of moving lines (17^th^ hour, period 97-102), did not induce an increase during the first 10 minutes or a decrease afterwards, suggesting that the moving lines only affect activity when the lines are novel.

**Fig 6.**
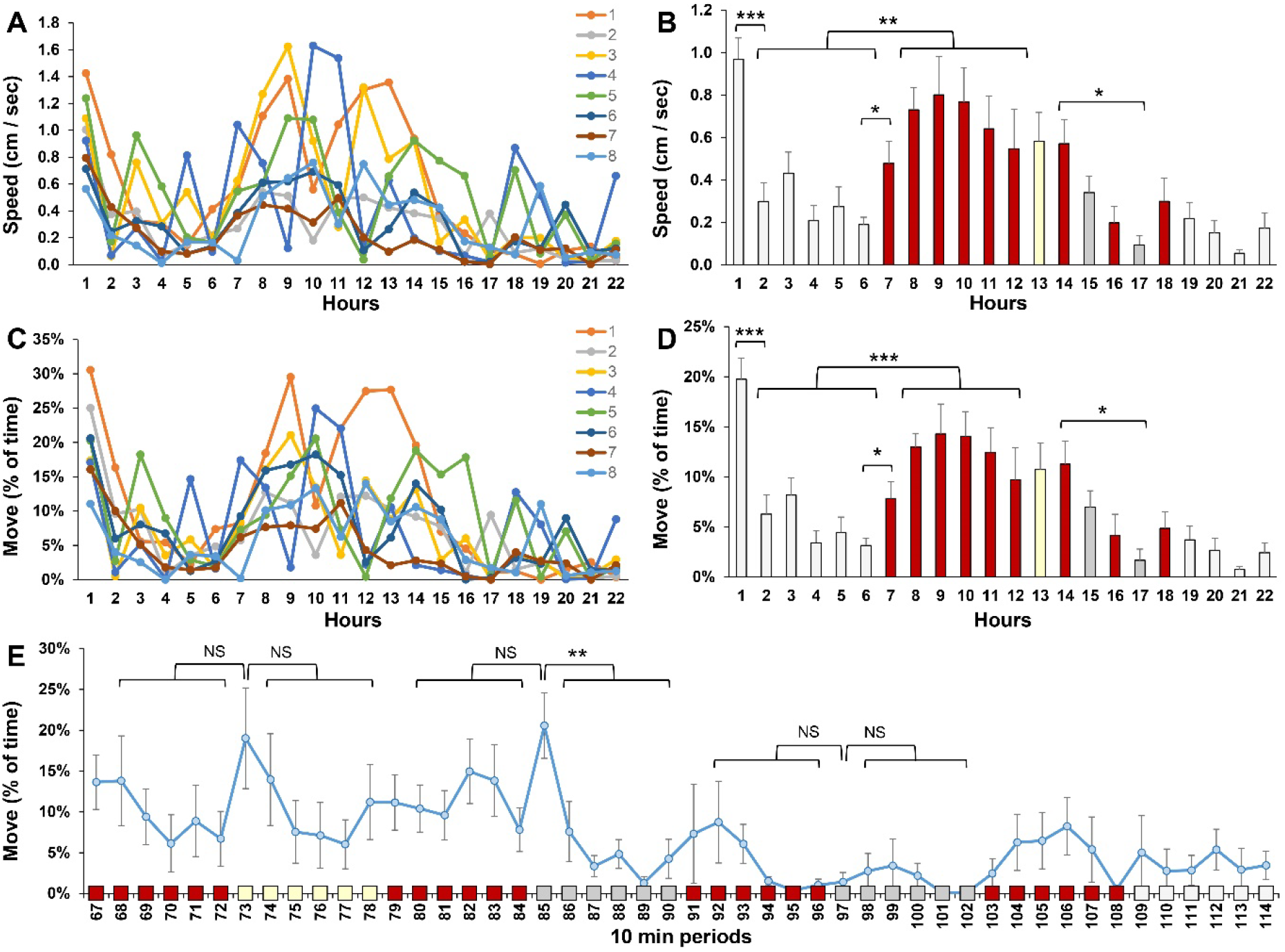
Activity of 8 mice in a 22-hour experiment. A) Speed of individual mice in a 22-hour recording. B) Average speed of 8 mice. C) Percentage of time that individual mice move in a 22-hour recording. D) Average movement of 8 mice). E) Movement in 10-minute periods (period 67-72 is the 12^th^ hour). * p<0.05, ** p<0.01, *** p<0.001, Welch test with a Bonferroni correction for multiple comparisons. Light gray bars = white background lighting. Red bars = red background lighting. Yellow bar = 13^th^ hour with a rotating yellow moth. Dark gray bars = 15th and 17th hour when the mouse house is illumined with bright light and moving red lines.

### Analysis of the stretch-attend posture

The stretch-attend posture (SAP) can be used as a measure of anxiety during risk assessment (*12*). SAPs are typically displayed when a mouse examines a novel object or novel environment. These mice do not move, or move slowly, and stretch their body towards the novel stimulus. In our analysis, we defined SAP as a movement below 1 cm/sec and an ellipse aspect ratio above 2.3. The ellipse aspect ratio of 2.3 corresponds to an ellipse eccentricity of 0.9, which was used in a previous study (*12*). SAPs were examined in a 22-hour recording of 8 mice (Fig 7), the same recording that was used to measure activity. We found that the mice displayed more frequent SAPs in the first hour (19%) than in the second hour of imaging (6%). The 13% difference between the first and second hour is significant (Welch test, p=0.007) and may be used as a measure of acclimation to the novel cage environment. SAPs were more frequently observed at nighttime than at daytime (17% vs. 6%, Welch test, p=0.01). Responses to visual stimuli were examined in 10-minute periods. The yellow moth did not induce changes in SAPs. The moving lines (period 85-90) induced more frequent SAPs during the first 10 minutes than the subsequent 50 minutes (15% vs. 3%, Welch test, p=3×10^−3^). The second set of moving lines did not induce significant changes in SAPs.

**Fig 7.**
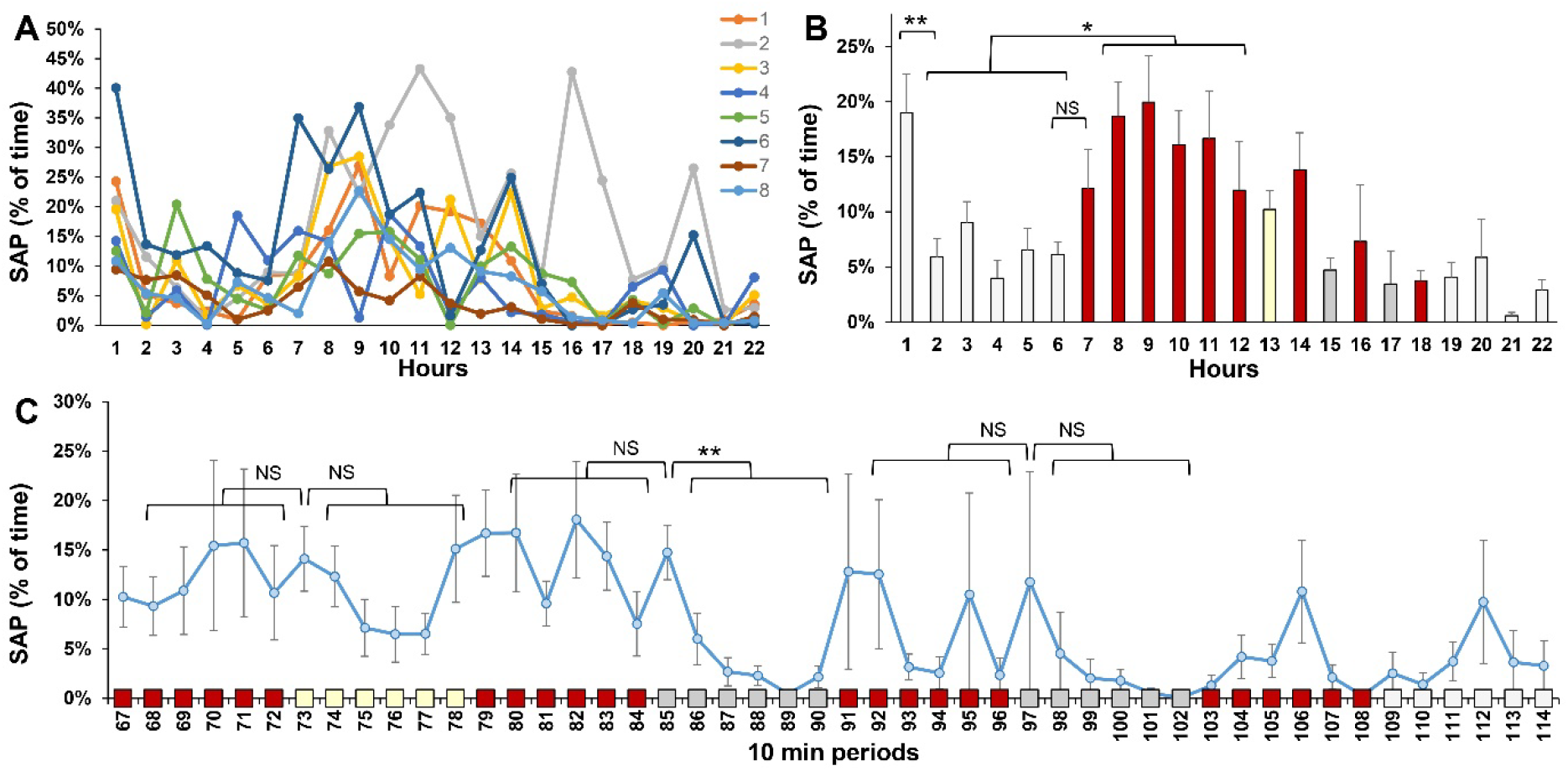
The Stretch-Attend Posture (SAP). A) The percentage of time that individual mice display SAP. B) Average SAP of 8 mice. C) Average SAP of 8 mice in 10 minute periods (period 67-72 is the 12th hour). * p<0.05, ** p<0.01, Welch test with a Bonferroni correction for multiple comparisons. Light gray bars = white background lighting. Red bars = red background lighting. Yellow bar = 13^th^ hour with a rotating yellow moth. Dark gray bars = 15th and 17th hour when the mouse house is illumined with bright light and moving red lines.

### Time in four quadrants

We examined the percentage of time that mice are located in each of the four quadrants of the cage (Fig 8). This percentage was compared to an expected 25% of time per quadrant in a random distribution. The mice displayed a variable preference for the home and window area, which did not reach statistical significance. The mice avoid the wall area during the first 11 hours of the imaging experiment, i.e. the mice spent less than 25% of their time in this quadrant (Welch test with a Bonferroni correction for 22 comparisons, p = 1×10^−6^, 2×10^−9^, 9×10^−9^, 3×10^−12^, 2×10^−11^, 6×10^−8^, 1×10^−8^, 2×10^−6^, 2×10^−4^, 4×10^−6^, and 3×10^−4^ for the first 11 hours of imaging). Similarly, the mice avoid the food area during the first 6 hours of the imaging experiment (Welch test with a Bonferroni correction for 22 comparisons, p = 7×10^−4^, 4×10^−4^, 4×10^−4^, 9×10^−7^, 2×10^−4^, and 6×10^−7^ for the first 6 hours of imaging). Based on these results, we conclude that mice avoid the wall area for 11 hours and the food area for 6 hours.

**Fig 8.**
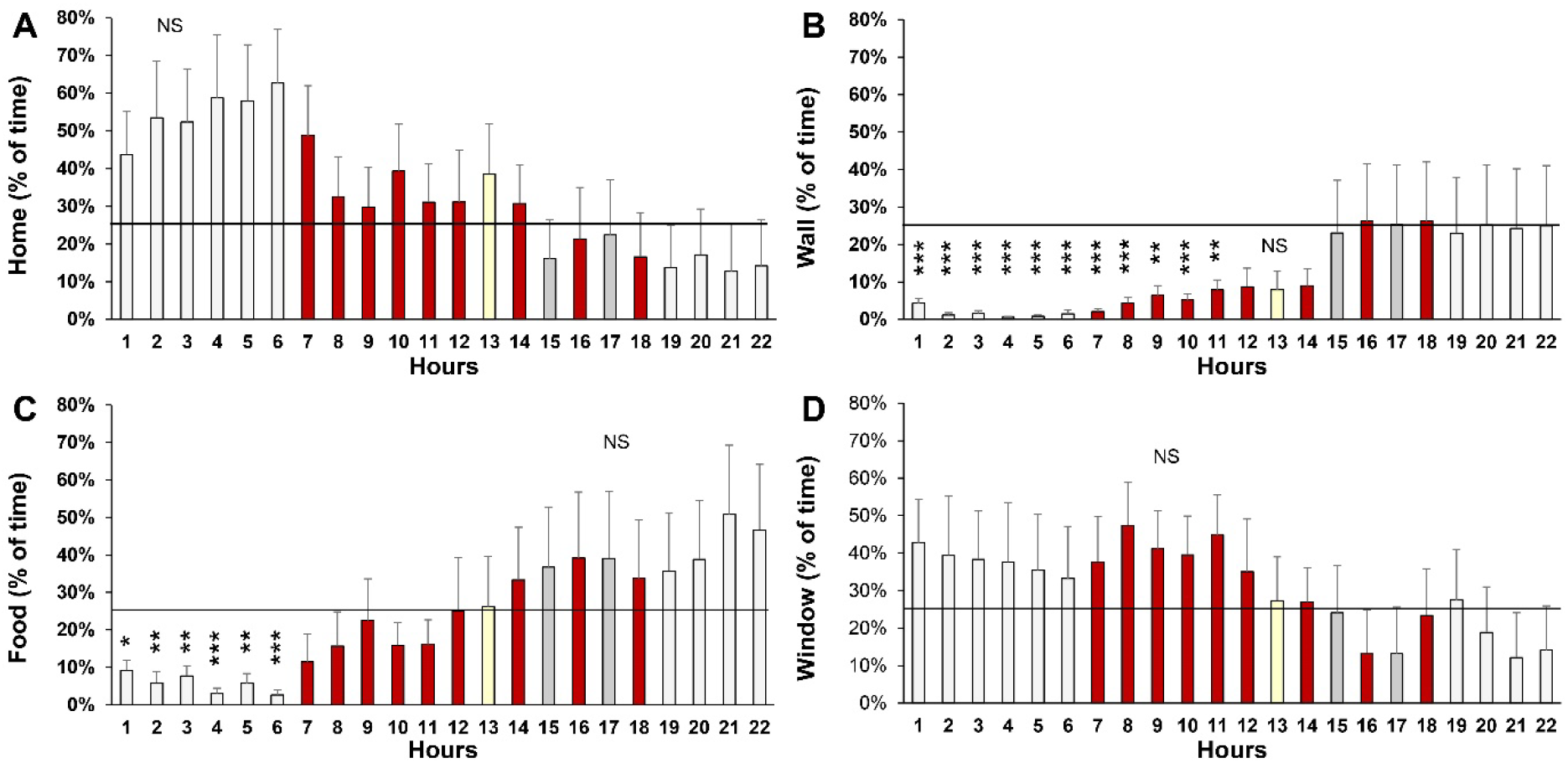
Time in four quadrants. A) Percentage of time in home area. B) Percentage of time in the wall area (an open area surrounded by two walls). C) Percentage of time in the food area. D) Percentage of time in the window area. * p<0.05, ** p<0.01, *** p<0.001, Welch test with a Bonferroni correction for multiple comparisons. Light gray bars = white background lighting. Red bars = red background lighting. Yellow bar = 13^th^ hour with a rotating yellow moth. Dark gray bars = 15th and 17th hour when the mouse house is illumined with bright light and moving red lines.

### Behavioral profiles

Various behaviors were summarized in a behavioral profile. The color-coded profiles provide an overview of movement, acclimation, habituation, and location in individual mice and in all mice as a group (Fig 9). The profiles show that individual mice display variable behaviors. For example, the mouse in cage 8 displays very little movement and a strong preference for the home area. Interestingly, the mouse in cage 8 was the only barber mouse, which had previously cut the whiskers of the mice in cage 7. The behavioral profiles can be used for analyses and comparisons between multiple imaging experiments, without creating oversized files. In addition, the profiles are suitable for hierarchical cluster analysis and may be used in future research to evaluate therapeutic treatments for various neural disorders (*13–15*).

**Fig 9.**
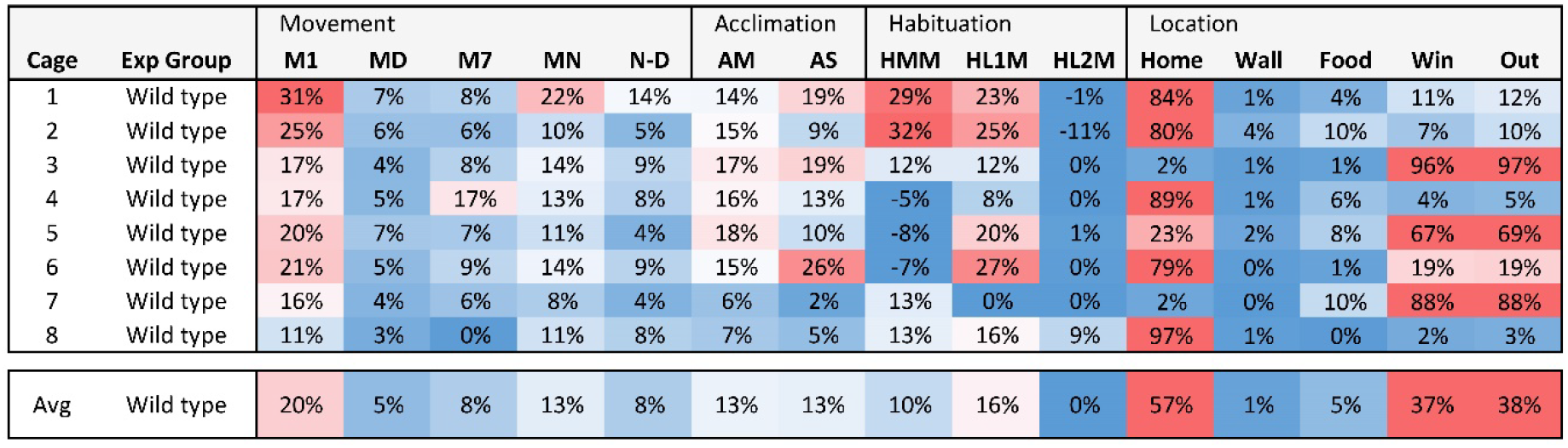
Behavioral profile. The behavioral profile summarizes 15 behaviors in 4-month-old female B6129SF2/J mice. These wild-type mice have a mixed B6 and 129S genetic background. In the F2 generation, each mouse is genetically unique. M1 = movement during 1st hour, MD = movement during daytime, M7 = movement in the 7th hour, MN = movement during nighttime, N-D = The difference between nighttime and daytime movement, AM = Acclimation to cage in % moved, AS = acclimation to cage in % SAP, HMM = habituation to a moth in % moved, HL1M = habituation to 1st set of moving lines in % moved, HL2M = habituation to 2nd set of moving lines in % moved. Home, Wall, Food, Win = % of time that a mouse is located in the quadrant with the red house, the empty quadrant surrounded by two walls, the quadrant with the food and water tray, or the quadrant with the screened window. Out = the sum of Wall and Window. The summary table is color-coded in a gradient from blue (0%) to white (15%) and red (30%), to provide a visual overview of the behavioral profiles.

## Discussion

The current study describes the development of an 8-cage imaging system, with animated visual stimuli, for overnight recordings of mouse behavior. This imaging system is, to some extent, a larger version of a system that we previously built for zebrafish (*15, 16*). When building the 8-cage imaging system for mice, we aimed for a system that 1) has sufficient throughput to study transgenic mice, mutant mice, and pharmacological treatments, and 2) can be built and duplicated on a limited budget. The 8-cage imaging system is well-suited for imaging a large number of mice. When imaging on a daily basis, the 22-hour experiments leave 2 hours for moving mice and cleaning cages. Thus, 32 mice can be imaged per week (8 mice x 4 overnight recordings). The system can be duplicated on a limited budget if additional capacity is needed. The most expensive components of the system are the standard laptop computers to control the visual stimuli and acquire images. The image analysis programs (ImageJ, Fiji, and DeepLabCut) are freely available and the PowerPoint file with visual stimuli, ImageJ macro, and MS Excel template can be downloaded from the supplementary information (S1, S2, S3). Additional information on the analysis in DeepLabCut is provided on GitHub. We also aimed for an imaging system that is easy to use. The PowerPoint presentations with visual stimuli are easily adjusted for further assay development. The ImageJ analysis can be carried out on a standard desktop computer, using a point-and-click interface, and the data sets can be further analyzed in MS Excel. New users do not need to know any coding and quickly learn to operate the imaging system, analyze acquired data, and develop novel assays, with little training. Instead of using ImageJ or Fiji, it is also possible to carry out advanced analyses of mouse postures in DeepLabCut, which requires some experience with Python coding. We used Brown University’s super computer (OSCAR) for the analysis in DeepLabCut. However, such analyses can be carried out on a desktop computer.

The developed imaging system has the following limitations. 1) The system is not ideal for the analysis of behavior over several days. Bedding is needed for long experiments and such bedding interferes with the bottom illumination and the presentation of visual stimuli. 2) We did not include acoustic stimuli in our experiments. However, it would be relatively easy to add sound to the PowerPoint presentations and speakers in the imaging system (*15*). 3) To maintain manageable files for image analysis and data processing, we imaged behavior for 22 hours at 1 frame per second. Imaging at 30 frames per second should be possible but will produce massive data files that are not suitable for subsequent analysis in MS Excel.

Other imaging systems have previously been developed for long-term recordings in a home-cage environment. For example, the PhenoTyper developed by Noldus consists of an observation cage with a camera, LED lights, and various sensors and stimuli (*17*). This state-of-the-art system can be used for imaging mice for days on end in a home-cage environment with bedding, food, and water. While the throughput of a single PhenoTyper is limited, a well-funded laboratory or core facility can install multiple PhenoTyper systems for simultaneous imaging of individual mice (*18*). Alternatively, imaging methodologies have been developed to measure social behavior of groups of mice in a single cage (*17*). Home-cage analyses are typically focused on measuring spontaneous behaviors. However, it can be interesting to include specific behavioral tests in a home-cage environment.

Our 22-hour assay can be considered a ‘behavioral test battery’, a series of tests to obtain a comprehensive assessment of brain function. These test batteries typically start with the least stressful test and end with the most stressful test (*19*). We followed this principle starting with daytime and nighttime lighting without visual stimuli, then showing a novel object (rotating yellow moth), and finally showing a more aversive stimulus (a bright area with moving lines). This test battery was used to evaluate 15 specific behaviors (Fig 9), which provide measures of hyperactivity in a novel environment, daytime and nighttime activity, acclimation, habituation, and avoidance of specific locations. In a test battery, later tests may be affected by earlier tests, and such effects can be interesting. For example, the mice in our study showed a response to the first set of moving lines, but not the second set of moving lines. We propose that the mice quickly learn to ignore the first set of moving lines and remember these lines an hour later.

The behavioral profiles of individual mice can be averaged into a behavioral profile for a particular mouse line (Fig 9). In the current study, we show the behavioral profile of 4-month-old female wild-type mice with a mixed B6 and 129S genetic background. We look forward to carrying out additional studies with male mice, older mice, mutant mice, transgenic mice that carry human disease genes, and mice that received treatments for specific disorders. Ultimately, advances in the analysis of behavior will contribute to a better understanding of neural diseases and may lead to the identification of novel therapeutics.

## Methods

### Approval of animal experiments

All experiments were carried out in accordance with federal regulations and guidelines for the ethical and humane use of animals and have been approved by Brown University’s Institutional Animal Care and Use Committee (Animal Welfare Assurance Number D16-00183). We followed the PREPARE, 3R, and ARRIVE 2.0 guidelines to plan and carry out the experiments.

### Mice

One-month-old female B6129SF2/J mice were purchased from the Jackson Laboratory in Bar Harbor, Maine (JAX stock #101045). These wild-type mice have a mixed genetic background, originating from a cross between C57BL/6J females (B6) and 129S1/SvImJ males (129S). F1 offspring were crossed to generate F2 mice, i.e. the B6 129S F2/J from the Jackson Laboratory. The F2 mice are genetically unique and can have either black or agouti fur. We selected this line for two reasons: 1) assays developed in a mixed line are more likely to be widely applicable than assays developed in an inbred line, and 2) the B6129SF2/J mice are good controls for transgenic strains maintained on a mixed B6:129S background. The F2 mice were maintained in Brown University’s animal care facility in Optimice IVC racks on a 12-hour light / 12-hour dark cycle, with 4-6 mice per cage, and free access to standard mouse chow and drinking water. For optimization of the imaging configuration, 2 mice were imaged when they were 3-4 months old. These 2 mice were imaged in 5 separate imaging experiments. An additional 8 mice were imaged when they were 4 months old. These 8 naïve mice did not have any prior experience with the imaging system.

### The 8-cage imaging system

The imaging system was built inside a dark box (Fig 1). This dark box is 64 inches high, 39 inches wide, and 33 inches deep. A 36 x 24 x 1/2 inch transparent acrylic stage (Amazon, B00C13Z874) was mounted 30 inches from the floor. Two webcams (Logitech C922x Pro) were mounted on the ceiling of the box. The two cameras were connected to a laptop computer (Dell Latitude 5490).

A M5 LED pico projector (Aaxa Technologies) with a 900 lumens LED light source is used for background illumination and the projection of visual stimuli. This projector was mounted on the box’s front panel, with the projector lens 17 inches from the floor. The projector illuminates the bottom of the mouse cages via a mirror that lays flat on the floor. With the mirror, it is possible to illuminate all 8 cages within a relatively small box. The projector is connected to a second laptop computer with PowerPoint software. Visual stimuli are shown to the mice as PowerPoint presentations.

### Cages for imaging

We built 8 cages for imaging using white polypropylene cage bottoms from Thomas Scientific (Cat No. 1113H35, 11.5 x 7.5 x 5 inches, L, W, H). These cages were covered with 12 x 8 x 1/4 inch transparent acrylic sheets (Amazon, B00YV5M7EM). We drilled a 2.5-inch diameter hole in the cage and covered this hole with a screen (Everbilt, hardware cloth, 23-gauge wire, galvanized, 1/4-inch mesh opening) to provide a window for ventilation (Fig 1C). This window is in the lower right corner in cage 1, 3, 5 and 7, and in the lower left corner in cage 2, 4, 6 and 8. When the odd and even cages are placed next to each other, the mice can look at each other through these windows, but cannot touch each other due to the inward sloping walls of the cage. The cages also contained a transparent red mouse house (Bio-Serv, mouse arch, red) and a transparent 7-sided pill box for food and water (Amazon, B07TJ2GVRZ). The mouse house was bolted to the side of the cage and the pill box was bolted to the mouse house. Two square pieces of food (Teklad Global 14% Protein Rodent Maintenance Diet) were placed in the feeder with an empty section in between. The remaining sections were filled with water. The cages were cleaned with 70% ethanol at the end of an experiment.

### Nuts and bolts

For investigators interested in building their own imaging system, we include a few specs on the nuts and bolts. To hold the acrylic stage, we attached 5 x 3-inch heavy-duty shelf brackets (Amazon, B07HHFJNX2) to the inside of the dark box, attached two self-adhesive clear silicone bumper pads (Amazon, B08BR6CGWL) on top of each bracket, and floated the stage on these bumper pads. The projector was mounted on the front panel of the dark box using a Camolo z-mount bracket (Amazon, B09MQLX9QC). The screen, mouse house, and pillbox were attached in the cage using M4 25mm bolts (Amazon, B012TCNAD0), M4 10mm bolts (Amazon, B012TCNMJ2), M4 nuts (Amazon, B07H3VF3BF), and M4 washers (Amazon, B015A39K02). The transparent acrylic lid was attached to the cage using elastic strings that were fed through 8 small holes in the lid and wrapped around four M4 wingnuts (Amazon, B07QB5NKDG) on the outside of the cage.

### Behavioral assay

Mice were moved from the animal care facility to the imaging room 30 minutes before the onset of imaging. Individual mice were randomly transferred to the 8 imaging cages. Identifying information of individual mice was recorded and available to all researchers in the lab (not blinded). Mice were imaged for 22 hours, starting around 2 pm. We aimed to measure multiple behaviors during this time. Lighting is controlled using a PowerPoint presentation with automated slide transitions and animations. Slide 1 is used for the alignment of the cages in the imaging system. Slide 2 provides a light gray background for daytime illumination during the first 6 hours of the assay. Slide 3 provides a red background for nighttime imaging during the next 6 hours. Thus, nighttime in the imaging system starts around 8 pm, as it does in the animal care facility.

Since mice are nocturnal and do not have red photoreceptors, we hypothesized that the mice are relatively active during this period. Slides 4-9 are shown for 1 hour each and include various visual stimuli. Slide 4 shows a rotating yellow moth in the Wall area on a red background. Slide 5 provides a red background. Slide 6 illuminates the Home area with a light gray background and moving red lines. The other quadrants are red. Slide 7 provides a red background. Slide 8 again illuminates the Home area with a light gray background and moving red lines. Slide 9 provides a red background. The final slide, slide 10, provides a light gray illumination for the remaining 4 hours. The PowerPoint presentation is available in the Supplementary Information (S1).

### Image acquisition

The two webcams were controlled by SkyStudioPro (version 1.1.0.30), a freely-available program developed by DjSadhu in the Netherlands (skystudiopro.com). SkyStudioPro combines the two camera feeds into a single recording (Fig 2). We used the following camera settings, aiming for 1) consistent illumination in the camera’s red channel, 2) manageable files, and 3) under one million rows of data for the final analyses in MS Excel. To prevent white balancing at nighttime, the ‘Capture devices’ were set at a fixed white balance of 4000. In the Camera control, the focus was set at 0, exposure at −5, and the capture size of each camera at 640 x 480 pixels (combined image = 640 x 960 pixels). We briefly unplugged camera 1 to set these values for camera 2. In Saving & Recording, the live and time-lapse codec was set to ffdshow video codec. In the Time lapse tab, we set the snapshot interval at 1 sec, movie speed at 30 fps, and autosave at 80,000 frames. With these settings, the 22-hour recording is saved as a 9 GB AVI file, and can be played back within an hour at 30x speed.

### Image analysis in ImageJ or Fiji

We developed an ImageJ / Fiji macro for automated analyses of the AVI movies (Fig 3). ImageJ and Fiji are freely available online and the macro is included in the supplementary information (S2). This macro can be installed in ImageJ or Fiji as a plugin (Open ImageJ or Fiji, Plugins, Install). The macro guides the user to open a movie as a virtual stack and collects information on the imaging files. For example, it asks the user to move a slider to the 12-hour point of the movie, when the first visual stimuli appear. It then uses this information to calculate the exact image interval. The macro then takes the first image in the stack, separates the color channels, and uses the red channel for subsequent analyses. In the red channel, the mice are visible at daytime and nighttime, even when they hide in their house. Within each cage, the macro applies a threshold and carries out a particle analysis, which filters out small objects (e.g. food pellets). The particle analysis is set to measure various parameters of the mouse, including its centroid, bounding rectangle, and best-fitting ellipse. X and Y coordinates are logged in a ‘Results’ file. The macro repeats these steps for all images in the AVI file, generating a long list of X and Y coordinates. These coordinates are then used to calculate the following parameters: location of the head, cage number, hour of imaging, period of imaging (in 10-minute increments), if a mouse is located in its ‘Home’, ‘Wall’, ‘Food’, or ‘Window’ area, if a mouse is ‘Out’ (Window or Wall), speed (in cm / sec), how often the mouse moved more than 1 cm / sec (% move), the aspect ratio of the ellipse (major / minor axis), and SAP, the stretch-attend posture. The mouse received a SAP score if its speed was smaller than 1 cm/sec and its ellipse aspect ratio was larger than 2.3. The ellipse aspect ratio was calculated as the major / minor axis of the best-fitting ellipse around a mouse without the tail. The ellipse aspect ratio of 2.3 corresponds to an ellipse eccentricity of 0.9, which was used in a previous study (*12*). In total, the ‘Results’ file has approximately 17 million data points (~ 600,000 rows x 28 columns).

### Image analysis in DeepLabCut

We tried out DeepLabCut as an alternative method for image analysis. This open-source software package uses training sessions and artificial intelligence (AI) to recognize specific body parts in various animals, including mice (*9–11*). DeepLabCut 2.2.3 (DLC) was used to perform markerless position estimation of a total of 48 points of interest: 12 mouse body parts, 15 cage features, and 21 different stimuli. To train the model, 430 frames were extracted from 10 videos using k-means clustering. This method of frame selection provides a set of images with different mouse postures, background lighting, and visual stimuli. Each frame was labeled using the DLC interface, and the complete collection of frames was evaluated to include at least 2 frames representing each stimuli. The trained network was based on ResNet-50 and the batch size was set to one. The network was trained for 400,000 iterations. We ran multiple behavioral sessions using two mice for optimizing the imaging system and DLC model training. All training videos were originally captured in 640×960 format, featuring an 8-cage set-up. Each video was cropped using Python 3.7, OpenCV, ffmpeg. Areas of interest were recognized using the findContours method implemented in OpenCV. DLC determined X and Y values for each of the 48 points of interest, along with likelihood values ranging from 0 to 1. The data generated by DLC was subsequently processed using Python 3.7, Numpy, and Pandas. This additional data processing step allowed us to measure the following behavioral paradigms: Speed, Stretch Attend Posture (SAP), Time in Home, Percent Peek, as well as distance from: Food, Food Plate, Window, Moth, Worm, Cricket, Top Wall, and Bottom Wall. Low likelihood (< 0.90) data points were filtered out during this step. All our annotated images and videos, DeepLabCut models and post processing scripts are available on GitHub (https://github.com/Creton-Lab). This GitHub page will also be used to post future updates of supplement 1-4.

### Data analysis in MS Excel

We used MS Excel to summarize the 17 million data points of the ImageJ ‘Results’ file into more comprehensive tables and graphs. For the analysis, an ImageJ Results file is copied in the ‘Data’ sheet of a MS Excel template (Supplementary information, S3). This template then averages specific behaviors per cage and per hour (e.g. speed in cage 1, hour 1) and then averages these behaviors in all cages (the average speed of all mice in the first hour). These values are shown in graphs over time (22 hours). Responses to visual stimuli were examined in 10 minute periods (visual stimuli are first shown in period 73 after 12 hours of imaging). A ‘Summary’ table shows the following 15 behaviors in table format: 1) M1 = movement during 1st hour, 2) MD = movement during daytime (average of hours 2-6), 3) M7 = movement in the 7^th^ hour (1st hour of nighttime), 4) MN = movement during nighttime (average of hours 8-12), 5) N-D = The difference between nighttime and daytime (in % move), 6) AM = Acclimation to cage in % moved (difference of % moved during the 1^st^ and 2^nd^ hour), 7) AS = acclimation to cage in % SAP, the stretch-attend posture (difference between the 1^st^ and 2^nd^ hour), 8) HMM = habituation to a moth in % moved (period 73 vs. the next 5 periods), 9) HL1M = habituation to 1^st^ set of moving lines in % moved (period 85 vs. the next 5 periods), 10) HL2M = habituation to 2^nd^ set of moving lines in % moved (period 97 vs next 5 periods), 11) Home = % of time that a mouse is located in the quadrant with the red house (average of hours 2-6), 12) Wall = % of time that a mouse is located in the quadrant with walls (average of hours 2-6), 13) Food = % of time that a mouse is located in the quadrant with the food and water tray (average of hours 2-6), 14) Win = % of time that a mouse is located in the quadrant with the screened window (average of hours 2-6), and 15) Out = the sum of Wall and Window. The summary table is color-coded from blue (0%) to white (15%) and red (30%), to provide a visual overview of the behavioral profiles.

### Statistics

Statistical analyses were carried out in MS Excel 2016 using Welch’s test, an unequal variances t-test. This test is well suited for the continuous data in our studies and is recommended over Student’s t-test and the non-parametric Mann–Whitney U test when the distribution is heavy-tailed and variances are unequal between groups (*20*). A Bonferroni correction was applied for multiple comparisons. This conservative correction reduces the chance of false positives, which is important in assay development. The following Bonferroni corrections were applied. 1) For the effects of moving lines in the hourly analysis, we compared 3 groups (hour 14 vs.15, 16 vs.17, and 14 vs.17) and differences were considered significant when p < 0.017 (0.05/3). 2) For analyses of visual stimuli in 10 minute periods, we compared 2 groups (first period with visual stimuli vs. the average of the prior five and next five periods) and differences were considered significant when p < 0.025 (0.05/2). 3) For preference of location in one of the four quadrants, we compared 22 groups (22 hours vs. an expected 25% occupancy in a random distribution) and differences were considered significant when p < 2.3×10^−3^ (0.05/22).

## Supporting information

Supplement 1 - PowerPoint

Supplement 2 - ImageJ macro

Supplement 3 - Excel Template

Supplement 4 - Results

## Acknowledgments

This work was supported by the National Institutes of Health, grant R01 GM136906 (R.C.) and grant R01 GM136906 supplement (R.C., J.A.K.)

## Author contributions

T.D.R.H., N.R.J., S.V.G., J.A.K. and R.C. contributed to the methodology, investigation, data analysis, and review of the manuscript. T.D.R.H and R.C. wrote the original draft of the manuscript. J.A.K. and R.C. contributed to the study design and supervision of the project.

## Data availability

The data and computer codes are available in the supplementary materials and a GitHub site.

## Supplement

S1. PowerPoint presentation

S2. ImageJ macro

S3. Excel template

S4. Excel file with results

