## Supplement 1 - PowerPoint for "An 8-cage imaging system for automated analyses of mouse behavior"

### Slide 1
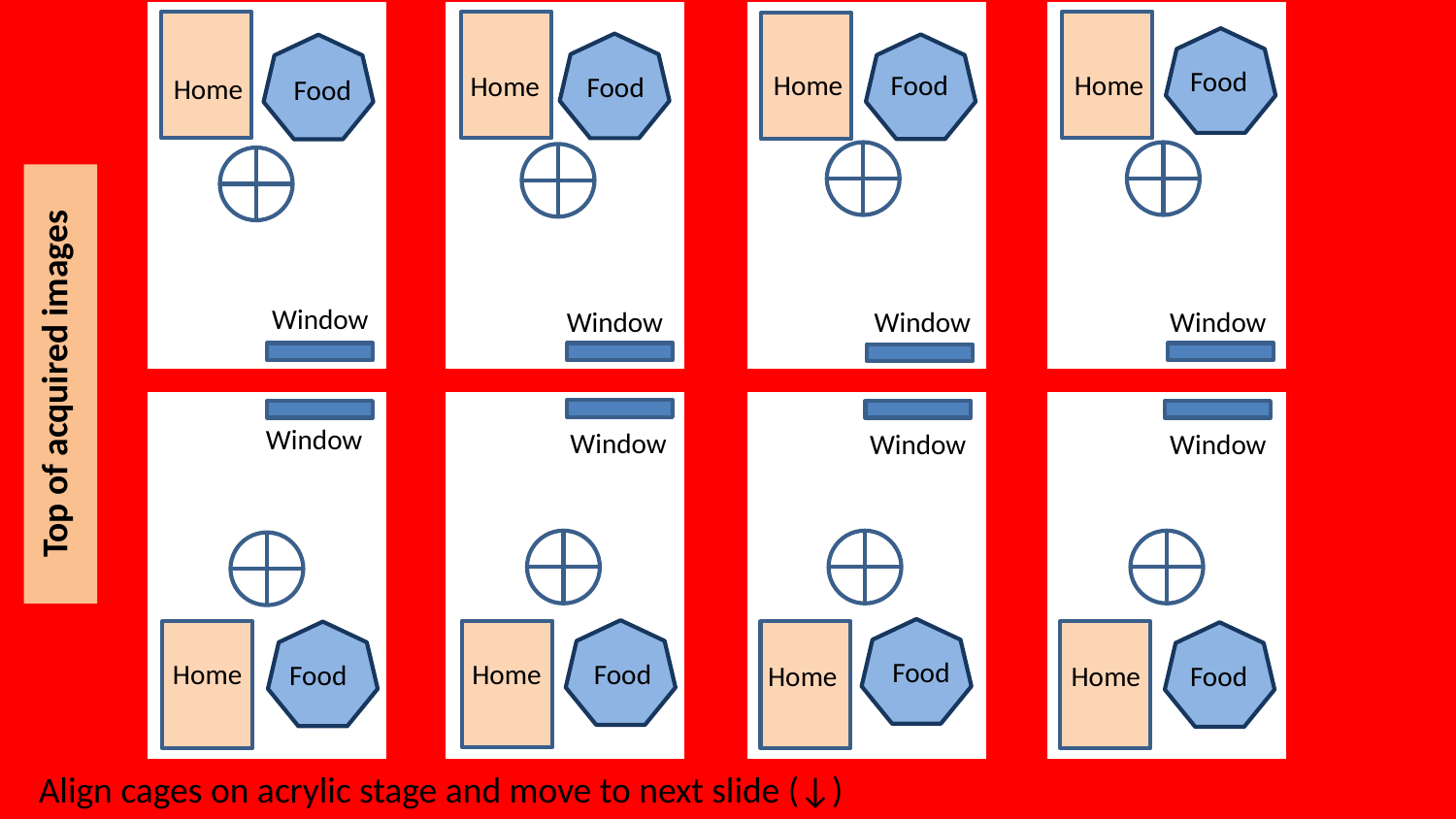

Food
Food
Home
Food
Home
Home
Food
Home
Food
,,
Top of acquired images
Window
Window
Window
Window
Window
Window
Window
Window
Food
Home
Home
Food
Food
Food
Home
Home
Align cages on acrylic stage and move to next slide (↓)

### Slide 2
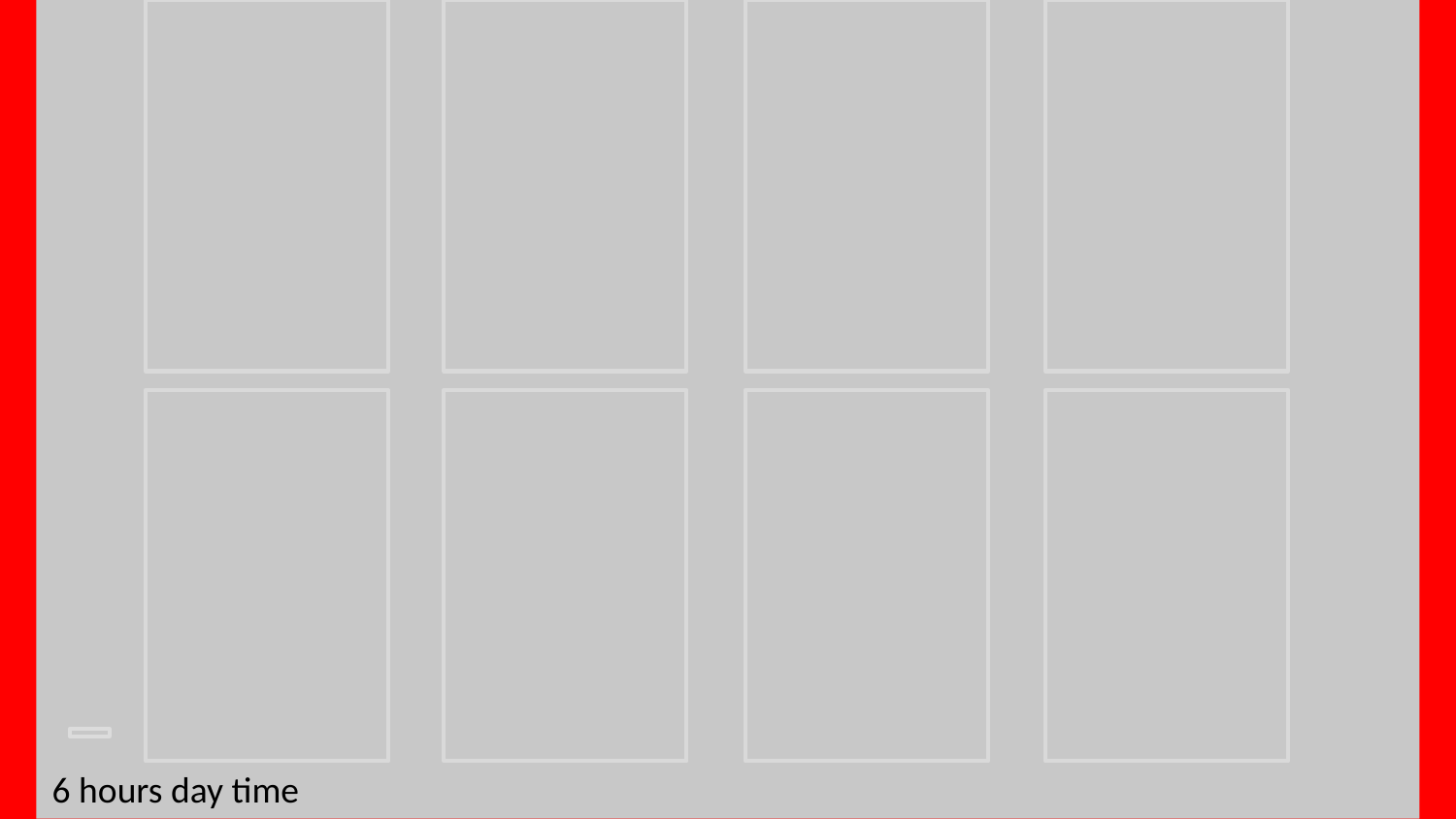

6 hours day time

### Slide 3
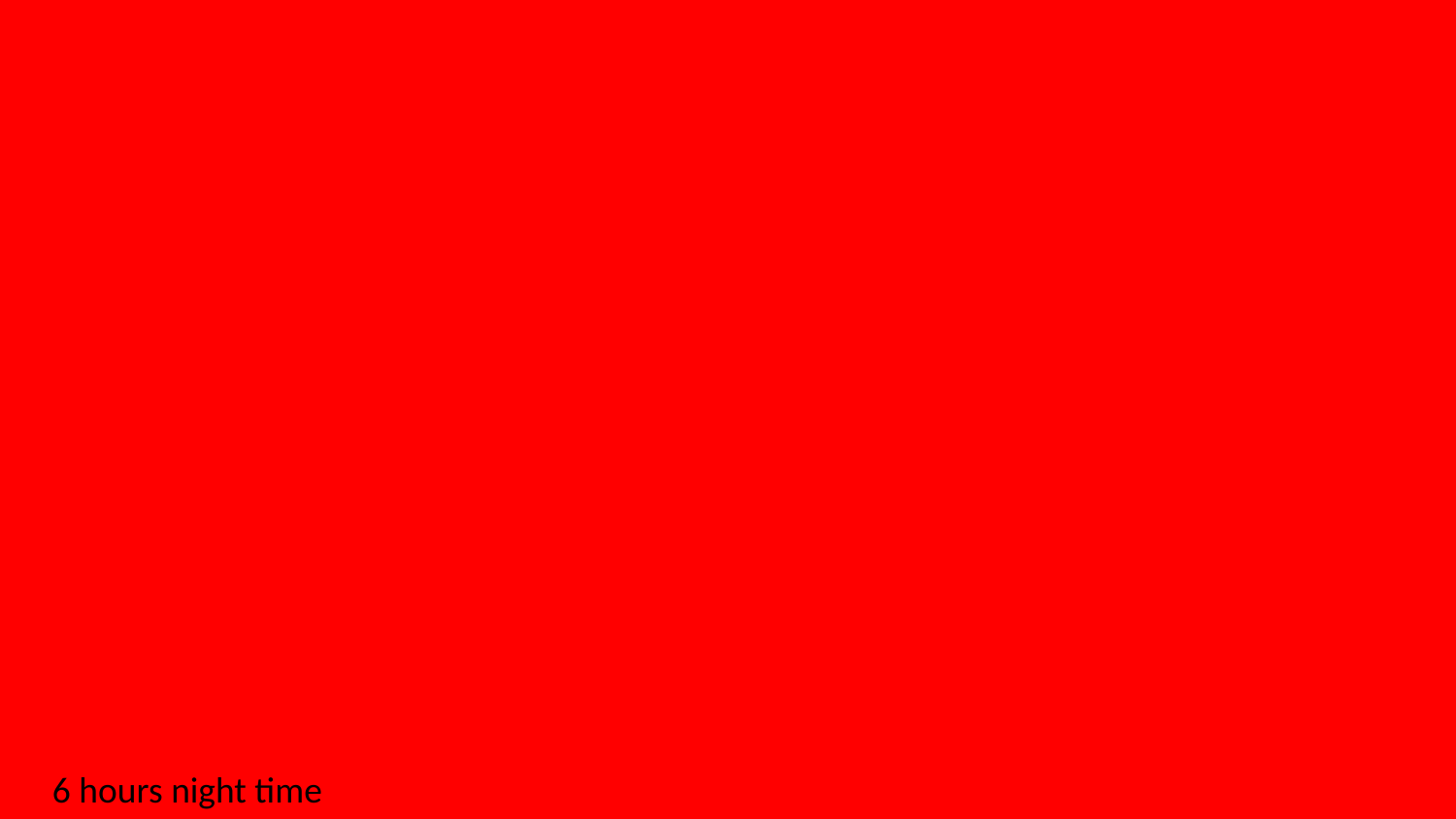

6 hours night time

### Slide 4
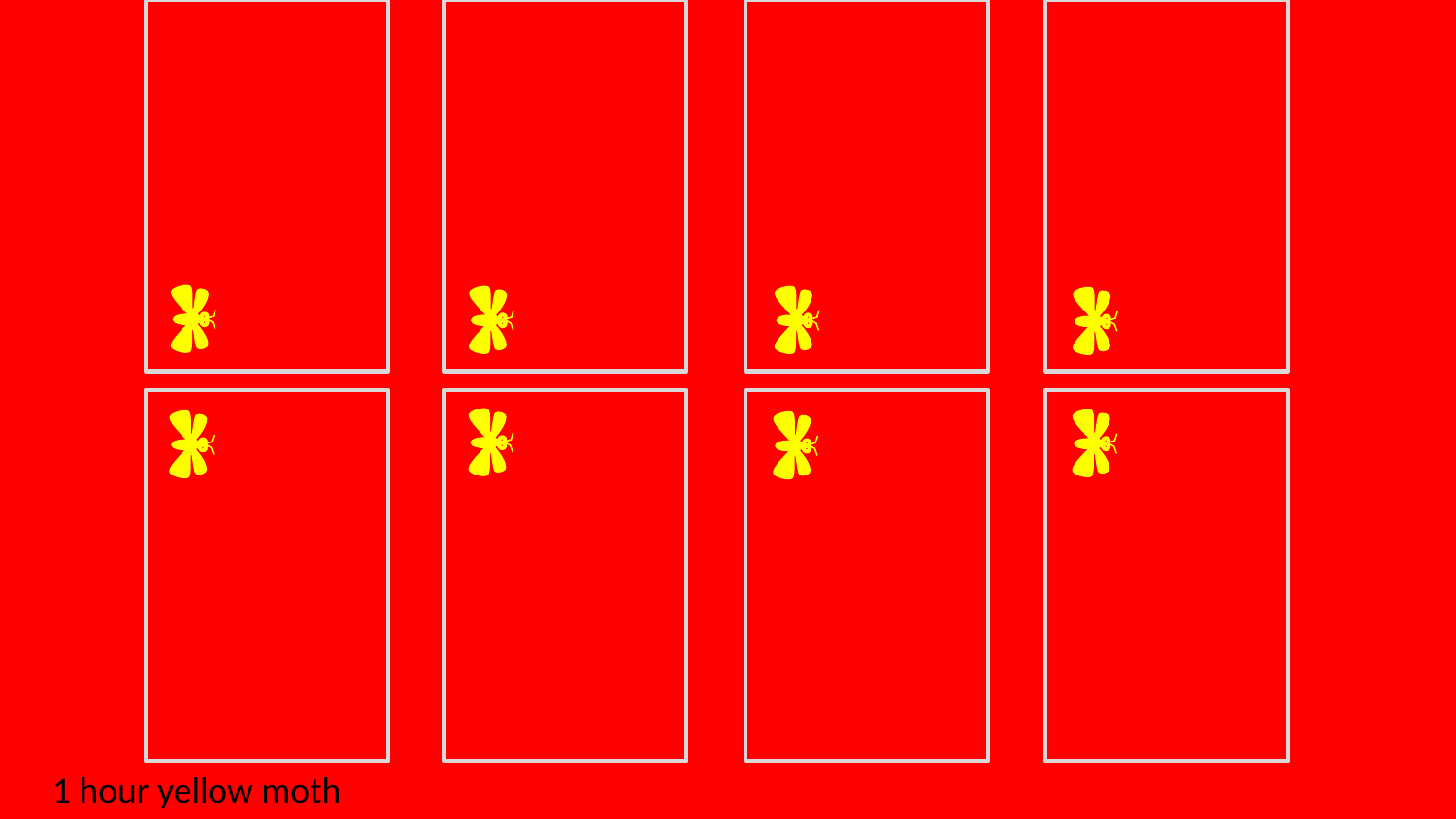

1 hour yellow moth

### Slide 5
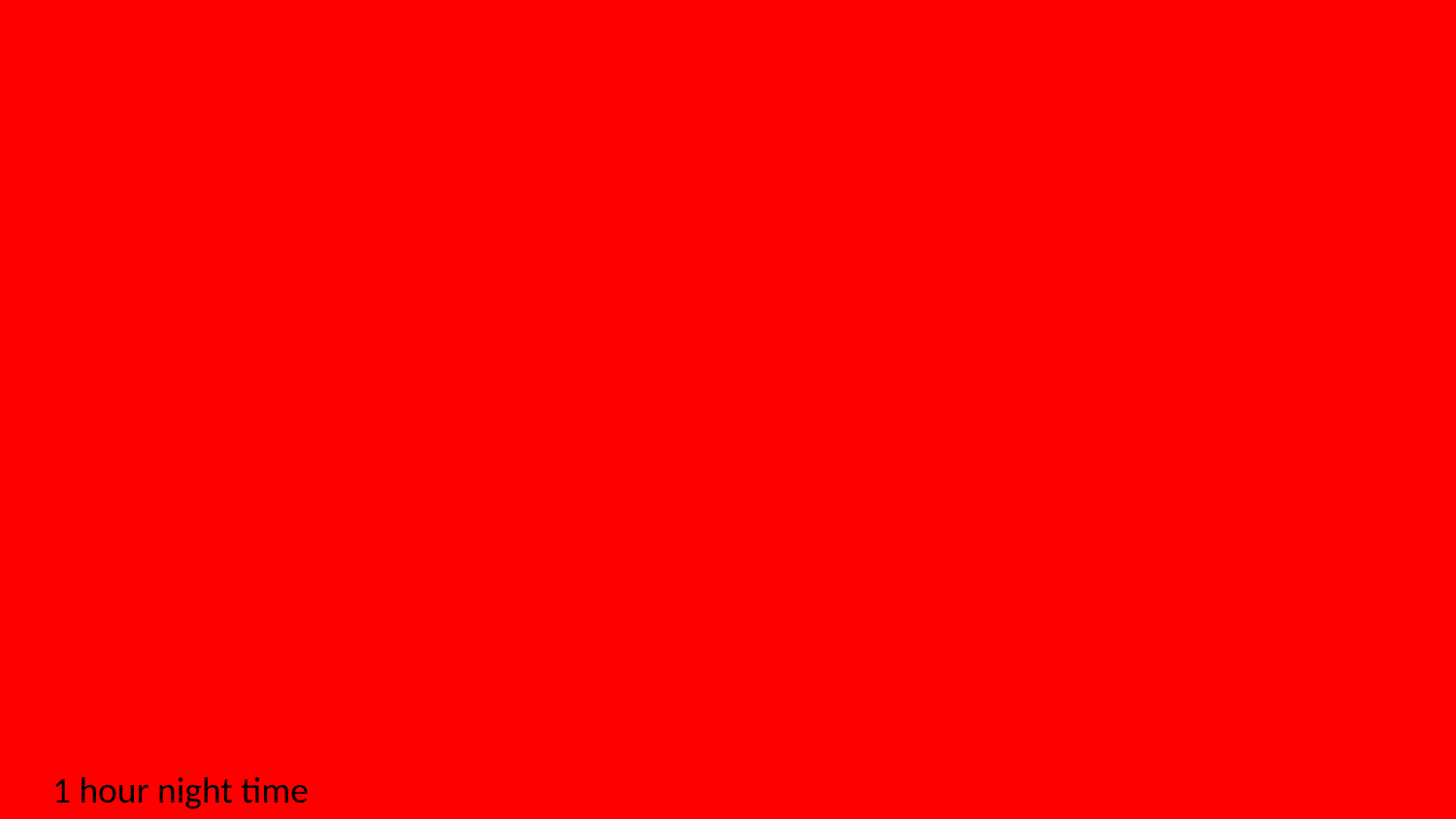

1 hour night time

### Slide 6
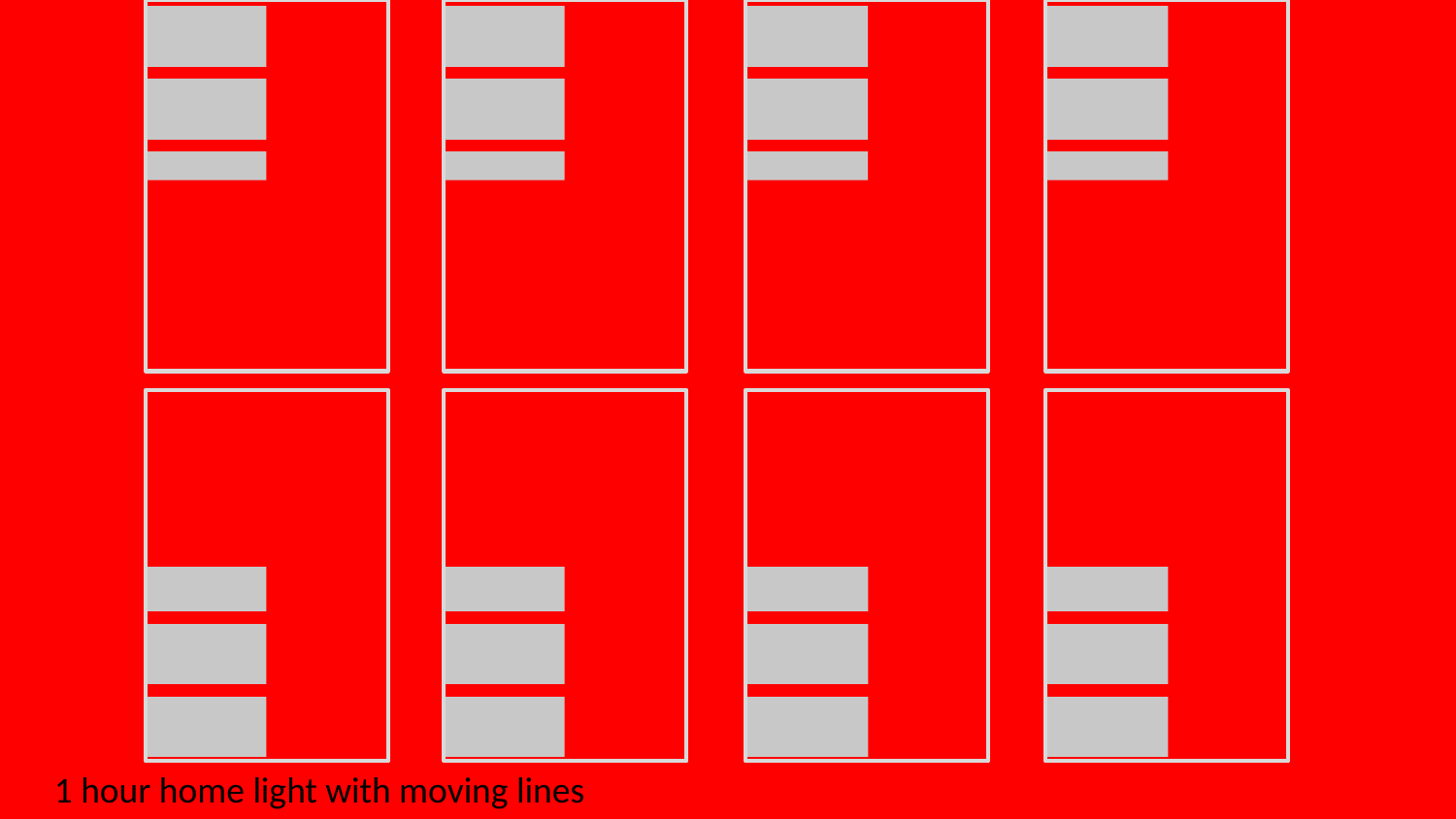

1 hour home light with moving lines

### Slide 7
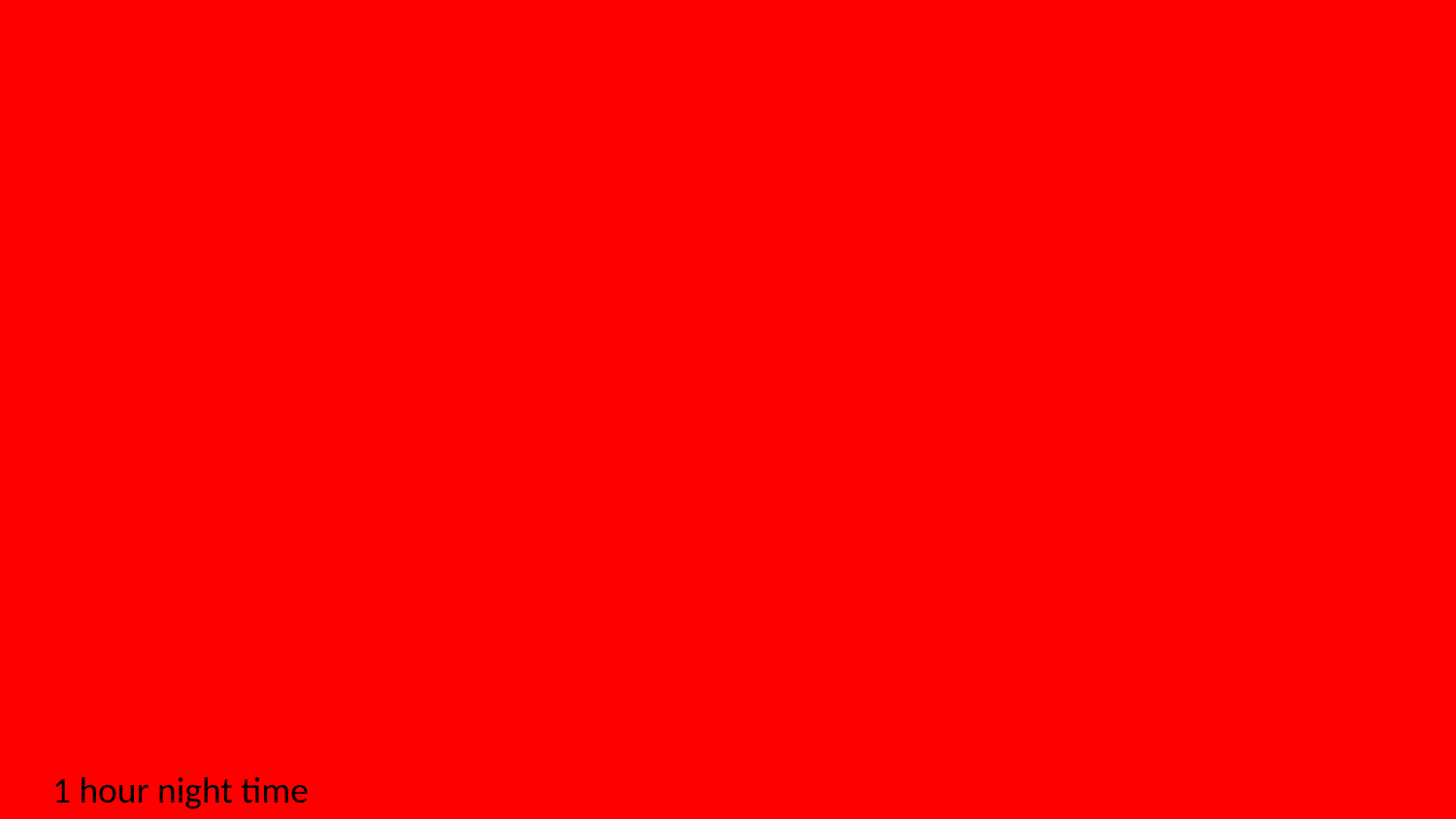

1 hour night time

### Slide 8
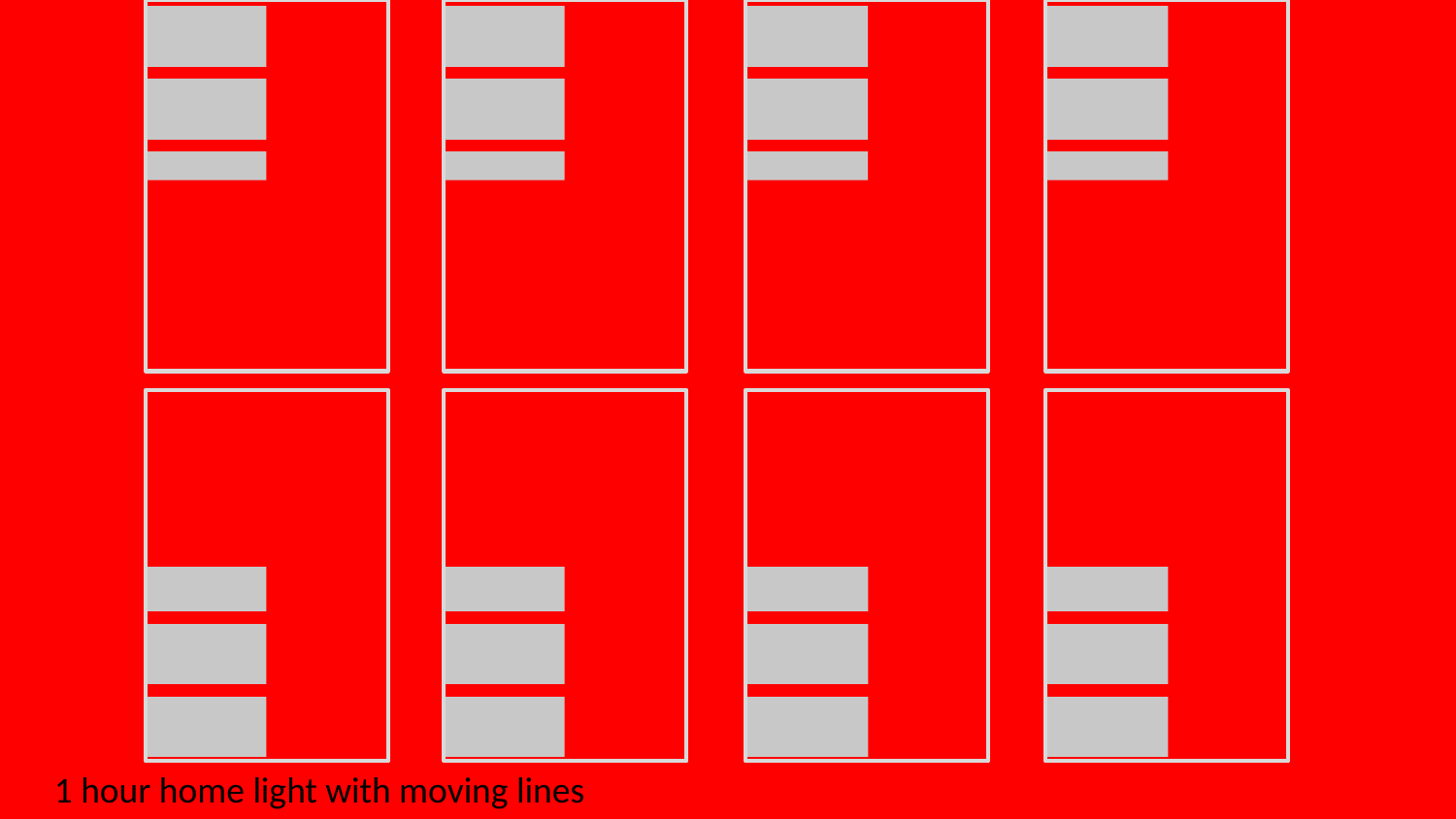

1 hour home light with moving lines

### Slide 9
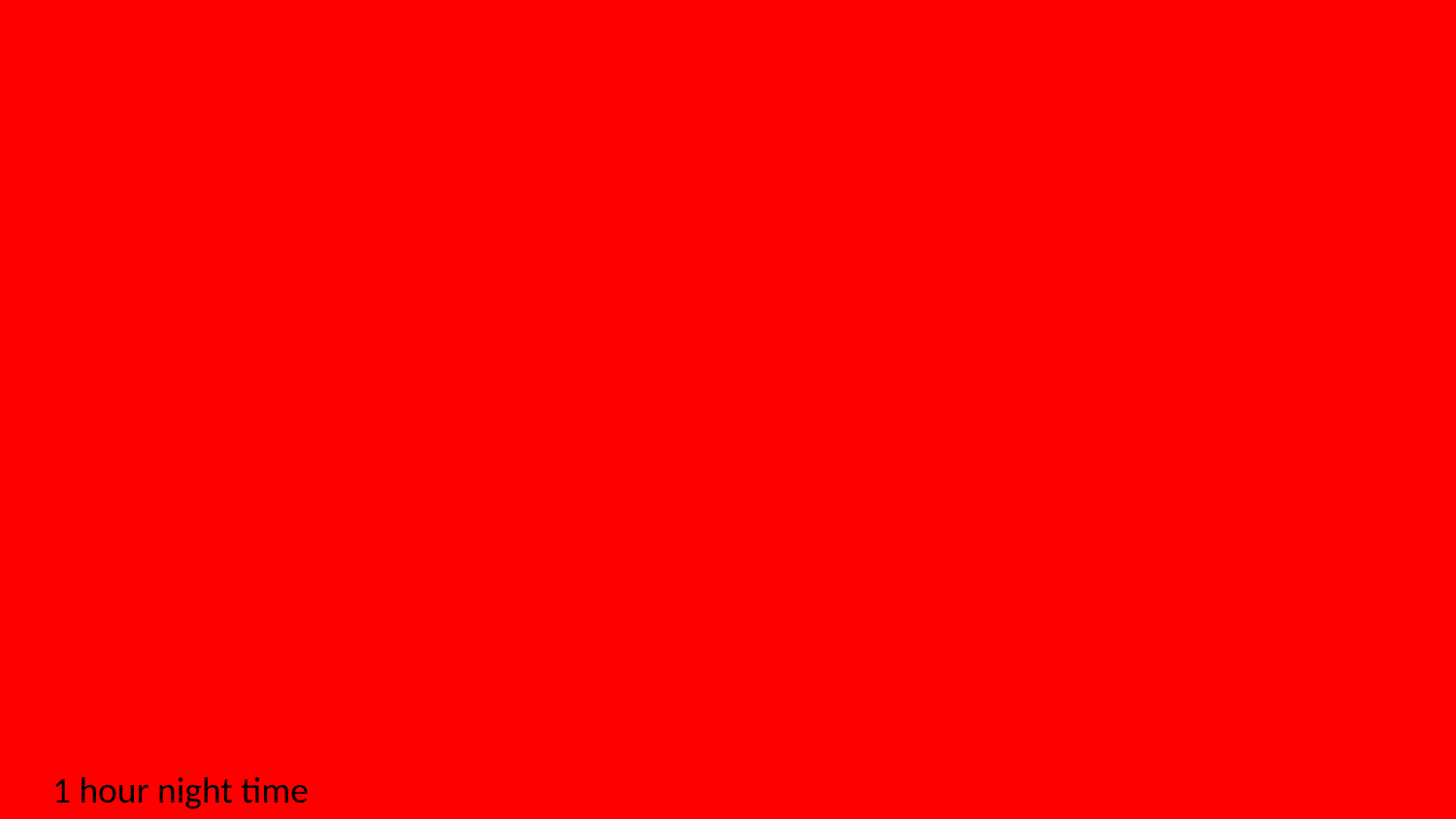

1 hour night time

### Slide 10
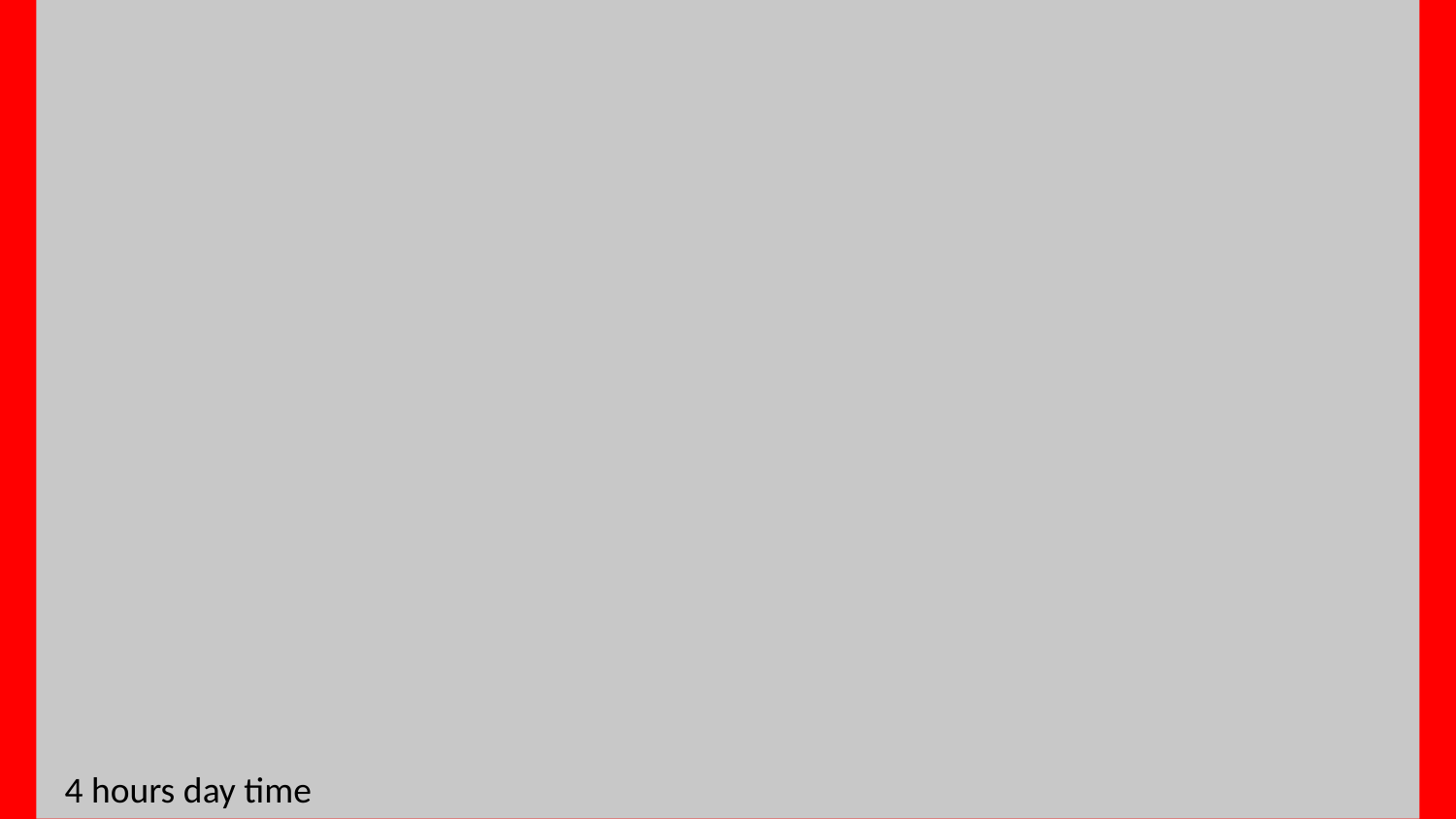

4 hours day time
